# Multiplex knockout of trichome-regulating MYB duplicates in hybrid poplar using a single gRNA

**DOI:** 10.1101/2021.09.09.459666

**Authors:** W. Patrick Bewg, Scott A. Harding, Nancy L. Engle, Brajesh N. Vaidya, Ran Zhou, Jacob Reeves, Thomas W. Horn, Nirmal Joshee, Jerry W. Jenkins, Shengqiang Shu, Kerrie W. Barry, Yuko Yoshinaga, Jane Grimwood, Robert J. Schmitz, Jeremy Schmutz, Timothy J. Tschaplinski, Chung-Jui Tsai

## Abstract

As the focus for CRISPR edited plants moves from proof-of-concept to real world applications, precise gene manipulation will increasingly require concurrent multiplex editing for polygenic traits. A common approach for editing across multiple sites is to design one gRNA per target; however, this complicates construct assembly and increases the possibility of off-target mutations. In this study, we utilized one gRNA to target *MYB186*, a known positive trichome regulator, as well as its paralogs *MYB138* and *MYB38* at a consensus site for mutagenesis in *Populus tremula* × *P. alba* INRA 717-1B4. Unexpected duplications of *MYB186* and *MYB138* resulted in a total of eight alleles for the three targeted genes in the hybrid poplar. Deep sequencing and PCR analyses confirmed editing across all eight targets in nearly all of the resultant glabrous mutants, ranging from small indels to large genomic dropouts, with no off-target activity detected at four potential sites. This highlights the effectiveness of a single gRNA targeting conserved exonic regions for multiplex editing. Additionally, cuticular wax and whole leaf analyses showed a complete absence of triterpenes in the trichomeless mutants, hinting at a previously undescribed role for the non-glandular trichomes of poplar.

**ONE SENTENCE SUMMARY:** Targeting conserved sequences with a single gRNA allowed efficient mutagenesis of a multigene family and the recovery of trichomeless and triterpene-free poplar mutants.

## INTRODUCTION

CRISPR (clustered regularly interspaced short palindromic repeats) technology has been adopted for plant genome editing in an increasing number of species for both basic and applied research (Bewg et al., 2018; Chen et al., 2019; Nasti and Voytas, 2021). The power of CRISPR is due in part to its simplicity with just two core components (Jinek et al., 2012): a nuclear-localized endonuclease, such as CRISPR-associated Cas9 that works universally across all domains of life and a synthetic guide RNA (gRNA) that is customizable and scalable for sequence-specific targeting. With its proven precision and efficiency (Li et al., 2013; Nekrasov et al., 2013; Shan et al., 2013; Endo et al., 2016) and given the polygenic nature of many agronomic traits, there is growing interest in targeting multiple loci for simultaneous CRISPR editing to aid gene function investigation and/or trait engineering (Armario Najera et al., 2019).

Multiplex editing usually involves coexpression of multiple guide RNAs (gRNAs). For the classic CRISPR/Cas9 system, this has been demonstrated using individual gRNA cassettes each driven by a separate RNA polymerase III (Pol III) promoter (Xing et al., 2014; Lowder et al., 2015; Ma et al., 2015). Alternatively, multiple gRNAs can be expressed in tandem with tRNAs as a single polycistronic transcript and processed into individual gRNAs using endogenous tRNA processing machinery (Xie et al., 2015). Polycistronic gRNA transcripts have also been engineered with built-in RNA cleavage sites for processing by ribozymes or the CRISPR-associated endoribonuclease Csy4 (Qi et al., 2012; Gao and Zhao, 2014; Tang et al., 2016; Čermák et al., 2017; Tang et al., 2019). In several cases, functional gRNAs were generated from a single transcriptional unit of *Cas9* fused with an artificial gRNA array without specific flanking sequences (Mikami et al., 2017; Wang et al., 2018). It has been reported that up to eight gRNAs have been successfully deployed for multiplex editing (Ma et al., 2015; Xie et al., 2015; Čermák et al., 2017).

An understudied approach is the use of single gRNAs to target homologous sequences at discrete loci. The capability was showcased by effective inactivation of all 62 copies of porcine endogenous retroviruses in an immortalized pig cell line using two gRNAs to target a highly conserved region of the polymerase (*pol*) gene (Yang et al., 2015). Besides parasitic elements, a single consensus gRNA has also been used to edit paralogs derived from various gene duplication events in soybean and sorghum (Jacobs et al., 2015; Li et al., 2018) or homoeologs in polyploid wheat and oilseed rape (Braatz et al., 2017; Zhang et al., 2017). Multiplex targeting of duplicated genes is especially important for investigation of functional redundancy in plant genomes that are shaped by whole-genome, segmental, tandem, and/or transposon-mediated duplications (Flagel and Wendel, 2009; Panchy et al., 2016). Depending on the duplication age and subsequent selection constraints, sequence similarity can be very high among duplicates, enabling identification of consensus target sites for multiplex editing by a single gRNA. This approach greatly simplifies construct design and assembly, reduces off-target potential that increases with the number of gRNAs (McCarty et al., 2020), and can be bundled with other multi-gRNA editing strategies discussed above for higher-order multiplex targeting of distinct gene families.

The present study explored the utility of a single gRNA for multiplex editing in an outcrossing woody perennial, *Populus tremula* × *P. alba* INRA 717-1B4 (hereon referred to as 717). As an interspecific hybrid, the 717 genome is highly heterozygous which presents additional challenges to gRNA design and edit outcome determination (Xue et al., 2015). Using trichomes as visual reporter, we targeted a known positive regulator of trichome development, *PtaMYB186* (Plett et al., 2010), and its close paralogs *PtaMYB138* and *PtaMYB38* for knockout (KO). We show that a single gRNA with SNP-aware design is effective for multiplex KO of paralogous genes and robust against copy number variations in a hybrid genome with an unexpected tandem duplication in one of its sub-genomes. We employed multiple approaches to address the analytical challenge of discriminating among highly similar target sites to discern mutations that ranged from small indels to large genomic dropouts. Finally, analysis of the resultant trichomeless mutants revealed a complete absence of triterpenes, and implicated a role for poplar trichomes in triterpene accrual.

## RESULTS

### Multiplex CRISPR/Cas9 editing of trichome-regulating MYBs

The known positive regulator of trichome initiation *PtaMYB186* (Plett et al., 2010) corresponds to gene model Potri.008G089200 in the *P. trichocarpa* v3.1 genome. It belongs to clade 15 of the R2R3-MYB protein family tree (Wilkins et al., 2009), which is expanded in poplar and contains three additional members, *MYB138, MYB38* and *MYB83*, with as yet unclear functions. The four clade 15 members are derived from multiple duplication events, based on whole paranome *K*_*S*_ (synonymous distance) distribution and gene collinearity analyses using the wgd program (Zwaenepoel and Van de Peer, 2018). These include an ancient (gamma) whole genome duplication (*MYB186* and *MYB83, K*_*S*_ = 3.76), a Salicoid duplication (*MYB186*/*MYB138* and *MYB38, K*_*S*_ = 0.21-0.22), and a tandem duplication (*MYB186* and *MYB138, K*_*S*_ = 0.0001) (Figure 1). MYB186, MYB138 and MYB38 share higher levels (88-96%) of amino acid sequence similarity than with MYB83 (55-57%). To ascertain these MYB involvement in trichome development, we mined RNA-seq data from different stages of 717 leaf development. Transcript levels of *MYB186, MYB138* and *MYB38* were highest in newly emerged leaves (Leaf Plastochron Index LPI-1) when trichome initiation occurs (Plett et al., 2010), but quickly declined thereafter in expanding (LPI-5) and mature (LPI-15) leaves (Figure 1). In contrast, *MYB83* transcripts were detected throughout leaf maturation (Figure 1), weakening support for its potential involvement in trichome development.

**Figure 1.**
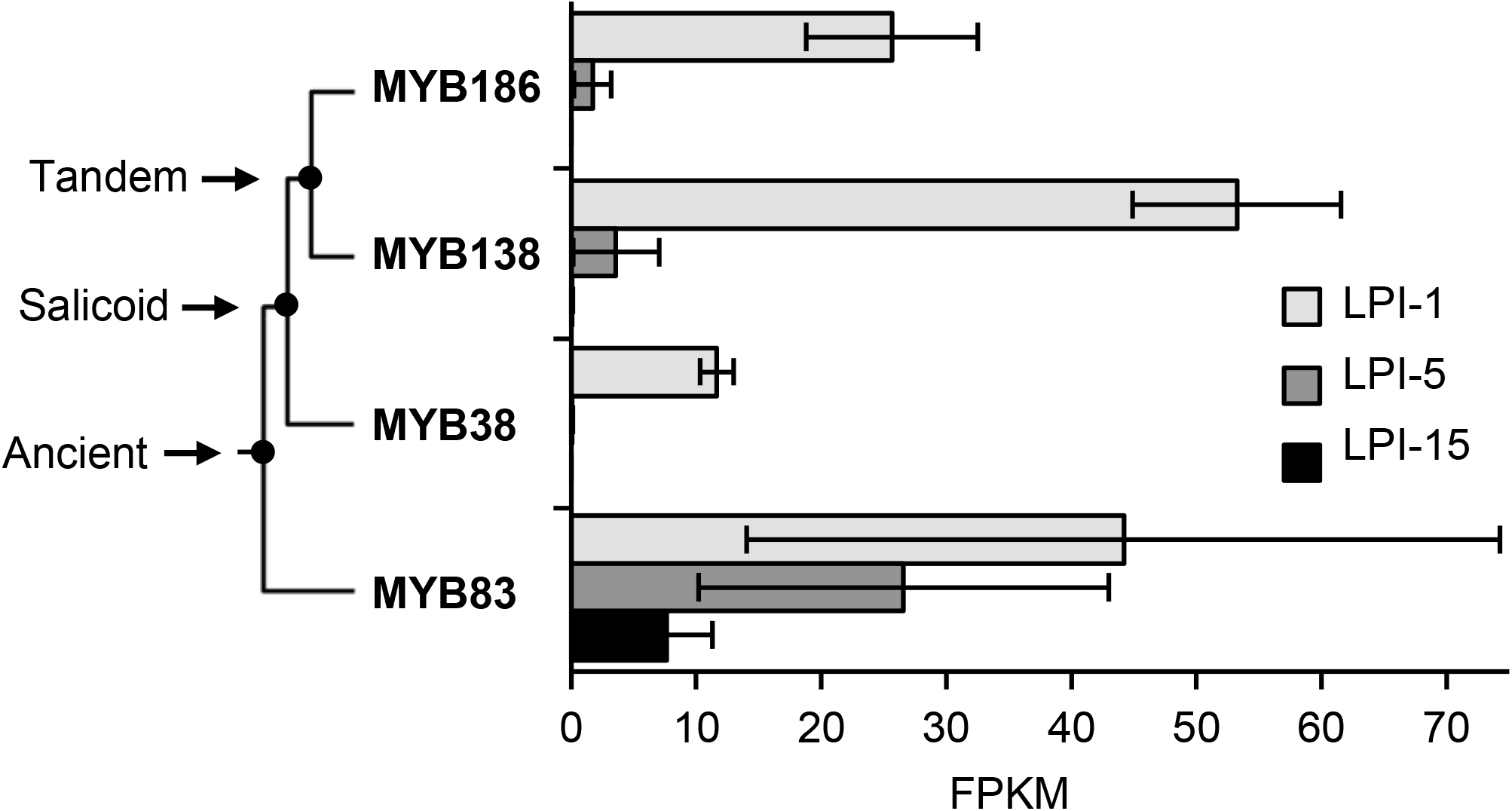
Expression of clade 15 *MYB* transcription factors during *Populus* leaf maturation. A simplified phylogenetic tree is shown with duplication history noted on the left. Data are mean±SD of n=3. LPI, leaf plastochron index; FPKM, fragments per kilobase of transcript per million mapped reads; MYB186, Potri.008G089200; MYB138, Potri.008G089700; MYB38, Potri.010G165700; and MYB83, Potri.017G086300.

We designed a single gRNA to target a conserved region in exon two of *MYB186, MYB138* and *MYB38* (Figure 2A) based on the *P. trichocarpa* v3.1 reference genome and cross-checked using the 717 variant database (Xue et al., 2015; Zhou et al., 2015) to assure the gRNA target sites were SNP-free in 717. Two CRISPR/Cas9 constructs were generated (see Methods); the first erroneously omitted a guanine between the gRNA and the scaffold sequences (referred to as ΔG, Figure 2B), which was corrected in the second construct (Figure 2A). Both constructs were used for 717 transformation in order to learn whether ΔG would affect CRISPR/Cas9 editing. In total, 28 independent events generated from the ΔG construct were all phenotypically indistinguishable from the wild type (WT) and Cas9-only controls (Figure 2C-J). In contrast, 37 independent events generated from the correct knock out (KO) construct were glabrous (Figure 2N-R), and one glabrous-like event (KO-27) had a greatly reduced trichome density across all shoot tissues (leaf, petiole and stem) independent of age (Figure 2K-M). SEM imaging revealed no trichome initiation or development on the abaxial leaf surface of the glabrous mutants (Figure 2Q). Epidermal cell morphology of young leaves from tissue cultured plants did not differ between control and mutant genotypes on either their abaxial (Figure 2F, N) or adaxial surfaces (Figure 2J, R). These results are consistent with roles for MYB186 (Plett et al., 2010) and its paralogs MYB138 and MYB38 in trichome initiation and development.

**Figure 2.**
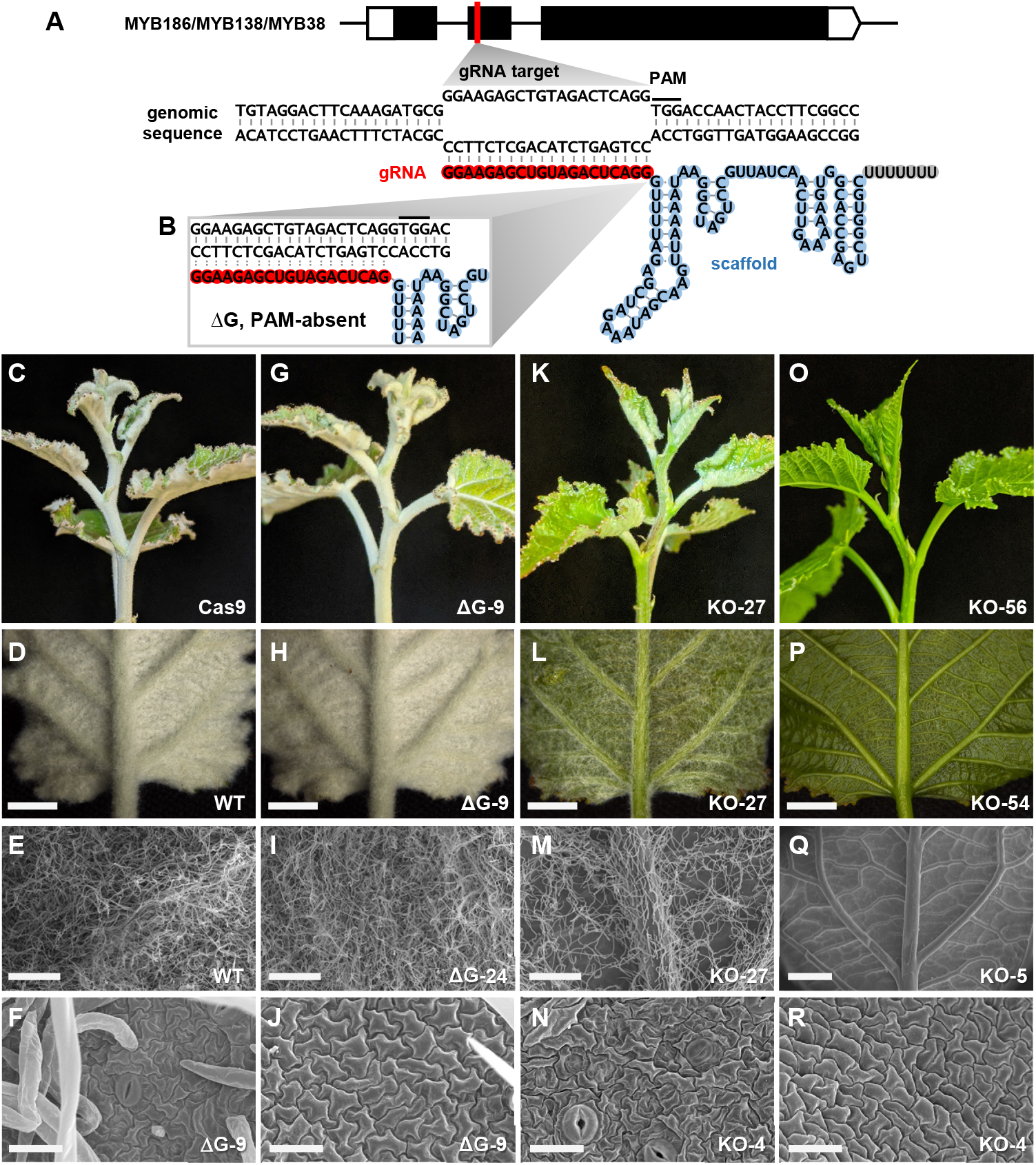
CRISPR/Cas9 KO of trichome-regulating *MYBs*. **A**, Schematic illustrations of the *MYB* gene structure, gRNA target site, and base pairing between the genomic target (black) and the gRNA spacer (red)-scaffold (blue) molecule. Black line denotes the protospacer adjacent motif (PAM). **B**, Zoomed-in view of the ΔG vector configuration at the gRNA spacer-scaffold junction with a guanine omission. **C-R**, Representative shoot tip (C, G, K, O) and LPI-1 abaxial (D, H, L, P) phenotypes and SEM images (E, F, I, J, M, N, Q, R) of soil-grown WT (D, E), Cas9 vector control (C), ΔG control (G-I), KO-27 (K-M), and null mutant (O-Q) plants, and leaf abaxial (F, N) or adaxial (J, R) images of tissue cultured ΔG (F, J) and null mutant (N, R) plants. Scale bar = 3 mm (D, H, L, P), 500 µm (E, I, M), 1 mm (Q), or 25 µm (F, J, N, R).

### Mutation spectrum of duplicated 717 _MYB_ alleles

A random selection of 30 glabrous events, 28 ΔG events, two Cas9-only events and four WT plants were subject to amplicon deep-sequencing using consensus primers for *MYB186, MYB138* and *MYB38*. Initial analysis by AGEseq (Xue and Tsai, 2015) showed numerous chimeric edits (mix of edited and unedited sequences at a given site) not observed in other CRISPR/Cas9-edited 717 transgenics in our experience (Zhou et al., 2015; Bewg et al., 2018; Tsai et al., 2020). *De novo* assembly of amplicon reads from control samples revealed seven distinct sequences, more than the expected six alleles of the three target genes. Blast search against the preliminary 717 genome assemblies by the Joint Genome Institute uncovered an unexpected copy number variation in 717 relative to the *P. trichocarpa* reference genome. The region containing paralogous *MYB186* and *MYB138* on Chromosome (Chr) 8 is found as a tandem duplicate in one of the 717 subgenomes (Figure 3A). This results in three alleles each for *MYB186* and *MYB138* (two on the main subgenome [Chr8m] and one on the alternative subgenome [Chr8a]) and two alleles for *MYB38* on Chr10 (Chr10m and Chr10a, Figure 3A). Two of the eight alleles were identical in the (original) amplicon region, explaining the seven distinct sequences we recovered from *de novo* assembly. Based on the 717 assemblies, we redesigned primers to ensure the amplicons span allele-specific SNP(s) to aid mutation pattern determination of the eight alleles.

**Figure 3.**
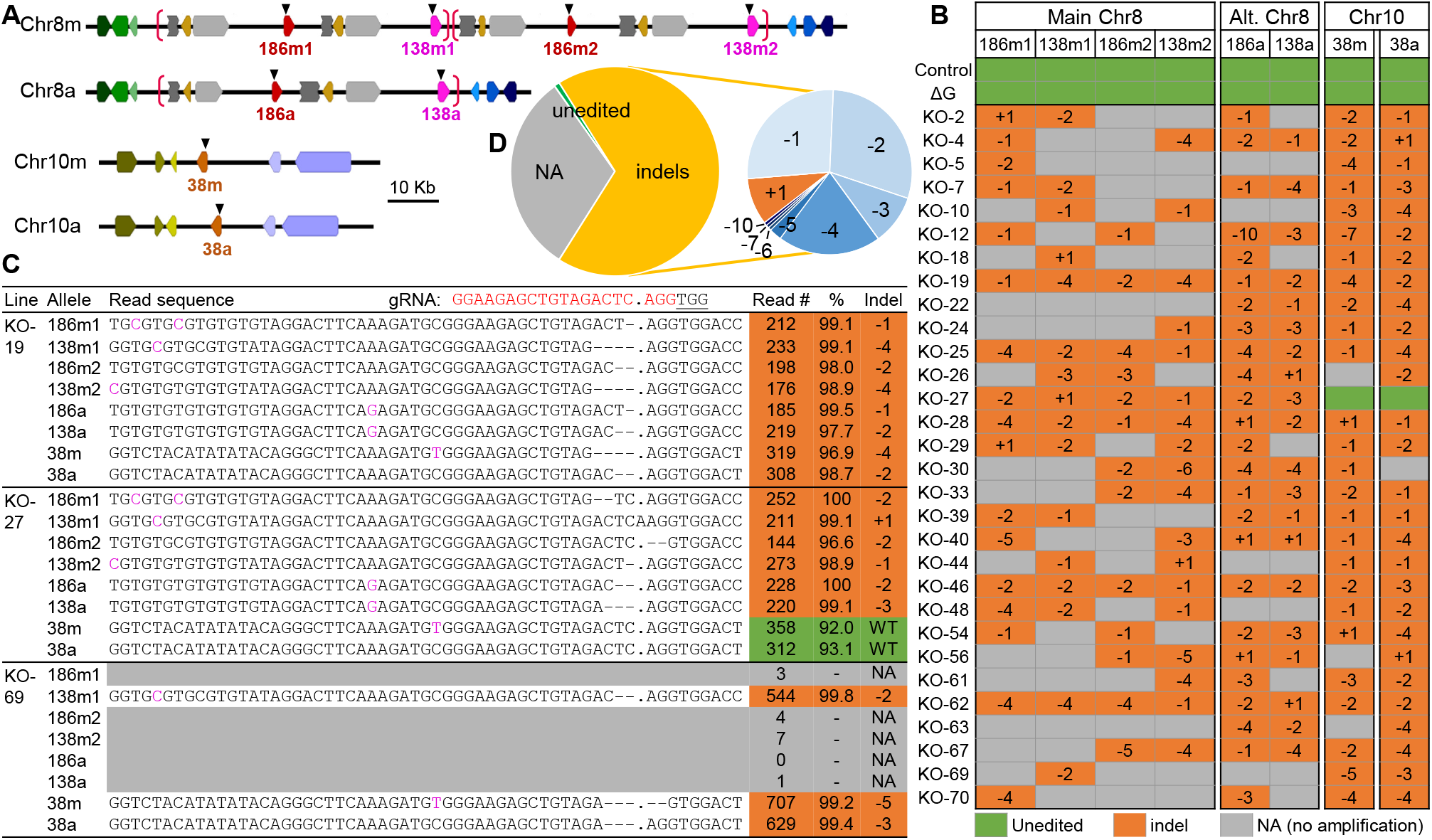
Mutation analysis of trichomeless mutants. **A**, Schematic illustration of *MYB186* and *MYB138* on Chr8 subgenomes (main and alternative, or Chr8m and Chr8a, respectively) and *MYB38* on Chr10m and Chr10a of the 717 genome. Neighboring genes are color coded for synteny and the putative duplication block containing *MYB186* and *MYB138* on Chr8 is marked by red brackets. Black triangles denote the eight gRNA target sites. **B**, Mutation spectrum determined by amplicon sequencing. The eight alleles are arranged by genomic position for each plant line and color-coded for the editing outcomes: green, unedited; orange, indel; and grey, no amplification (NA). **C**, Representative amplicon sequencing output of three mutant events. All eight alleles, their detection frequencies and indel patterns (mapped read count and percentage with the indicated pattern) are shown and colored as in B. The gRNA target sequence is shown on top and protospacer adjacent motif underlined. Allele-discriminating SNPs are shown in pink (see Supplemental Dataset S1 for the full data). **D**, Pie chart summary of the overall (left) and indel (right) editing patterns.

Amplicon-sequencing showed no editing in the 28 ΔG events, except one (ΔG-24) with a 9 bp deletion at one of the eight target sites (Supplemental Dataset S1). This translates into a mutation rate of 0.45% (one out of 224 potential target sites), which suggests a negative effect of the ΔG on CRISPR/Cas9 function (hereafter, the ΔG plants were treated as transformation controls). In contrast, we confirmed successful editing across the eight alleles in all glabrous mutants except KO-27 (Figure 3B-C, Supplemental Dataset S1). This event showed six edited and two WT (unedited) alleles, consistent with trichome detection in this line (Figure 2K-M). In aggregate, small insertions and deletions (indels) were the predominant edits at all sites (Figure 3B-D), with frameshift deletions of 1 bp (−1), 2 bp (−2) and 4 bp (−4) accounting for over three quarters of the indel mutations (Figure 3D). In-frame deletions (−3 or -6) accounted for 10% of indels and were detected in 14 events, including KO-27 (Figure 3B-D These in-frame mutations are unlikely functional because the gRNA target site is located within the third α-helix of the R2 domain critical for MYB-DNA interaction (Wang et al., 2020), and because 13 of the events with in-frame mutations are glabrous. We therefore conclude that all small indels we detected are null mutations.

### Large genomic dropouts between tandem genes

The vast majority (80%) of the sequenced mutants also harbored potentially large deletions as evidenced by the dearth of mapped amplicon reads at the target sites, referred to as no-amplification (NA) alleles (Figure 3B-D, Supplemental Dataset S1). The NA frequencies differed by chromosome position and were positively correlated with copy number, being highest at the Chr8m sites (four tandem copies), followed by the Chr8a sites (two tandem copies) and least at the single-copy Chr10 sites (Figure 3A-B). The NA alleles on Chr8 often spanned consecutive copies, suggesting large dropouts between two gRNA cleavage sites. To support this idea, we examined a subset of mutant lines using allele-specific primers for PCR amplification of the target genes. As expected, NA alleles yielded no PCR products, whereas alleles previously detected by amplicon sequencing produced observable PCR products (Supplemental Figure S1). We next used consensus primers for PCR amplification of all six Chr8 (*MYB186* and *MYB138*) alleles, approximately 850 – 950 bp, from three control plants and four KO lines each with 4-5 NA alleles on Chr8. These KO lines had reduced PCR band intensity when compared with controls (Figure 4A-B). Sanger sequencing of the PCR products resulted in clean chromatograms with clear nucleotide peaks throughout the sequenced length for KO-5 and KO-69 (Figure 4C), two mutant lines with only one detectable Chr8 allele (Figure 4B). In contrast, the chromatograms for KO-63, KO-70 (both containing two detectable Chr8 alleles) and WT samples were noisy as would be expected for mixed template (Figure 4B-C). The Sanger sequencing data of KO-5 and KO-69 not only confirmed the indel pattern (−2 in both cases) detected by amplicon sequencing, but also supported the occurrence of gene fusion between two gRNA cleavage sites, based on SNP patterns upstream and downstream of the gRNA target (Figure 4B-C). KO-5 harbors a fusion junction between *MYB186m1* and *MYB138m1* with a ∼29 Kb genomic dropout, whereas KO-69 contains a fusion of *MYB138m1* and *MYB138m2* with a ∼62 Kb genomic dropout (Figure 4B-C, Supplemental Figure S2). Both events likely contain additional large deletions or genomic fusions, as allele(s) downstream (KO-5) or upstream (KO-69) of the respective fusion point could not be PCR amplified (Figure 4B). Regardless, our findings show that a single gRNA is highly effective for multiplex KO of tandem duplicates via either small indels or large deletions.

**Figure 4.**
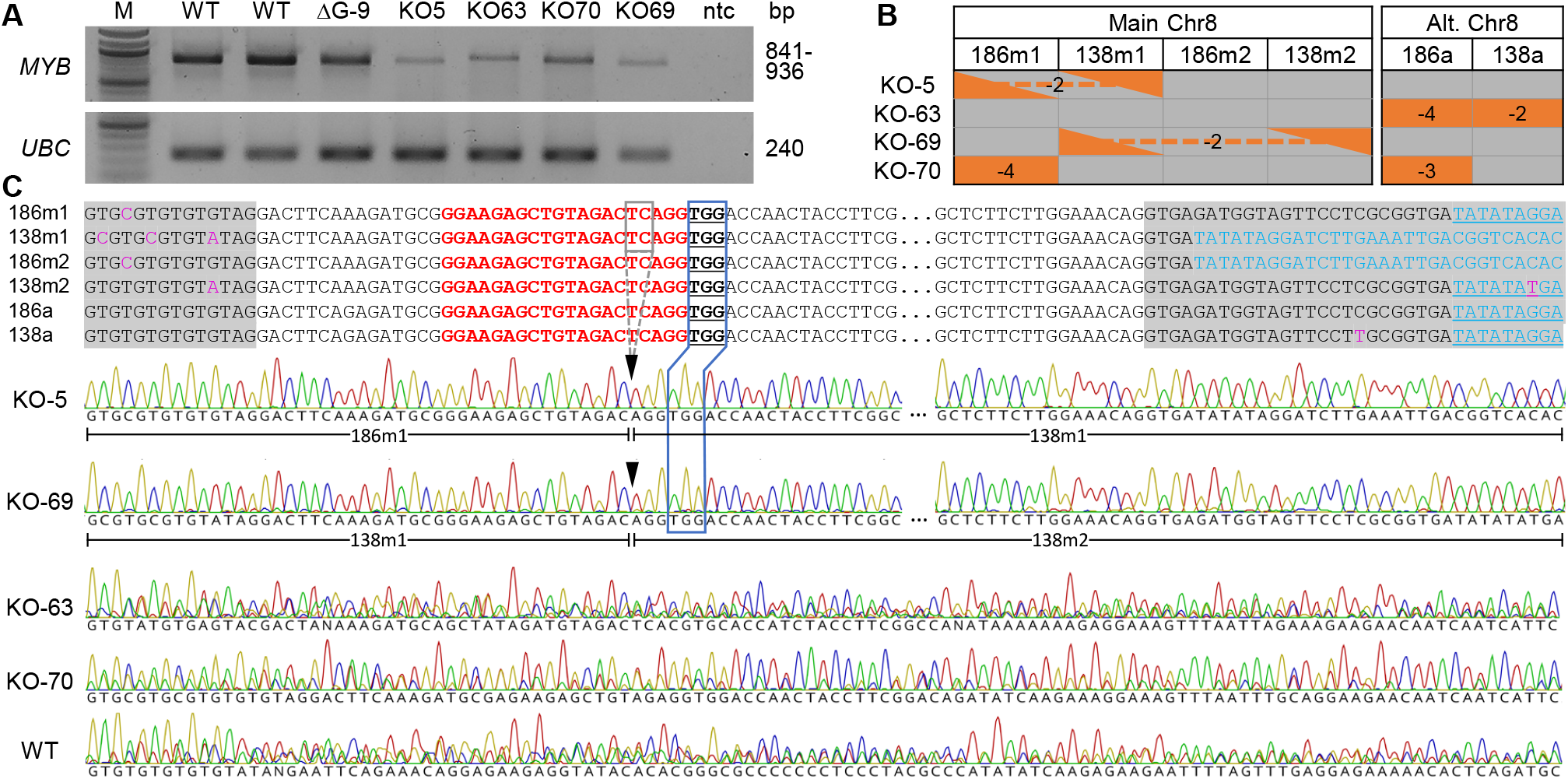
PCR analysis of selected mutant lines. **A**, PCR amplification of the six *MYB* alleles on Chr8 from two WT, one ΔG and four KO lines. The four KO lines were selected to represent one (KO-5 and KO-69) or two (KO-63 and KO-70) remaining Chr08 alleles. *UBC* (ubiquitin-conjugating enzyme) was included as loading control. M, molecular weight marker; ntc, no-template control. **B**, Mutation patterns of Chr08 alleles in the selected mutant lines as shown in Figure 3B. Orange triangles connected by dashes represent fusion junctions shown in **C**. C, Sanger sequencing of PCR products from A. Sequence alignment of the six alleles flanking the gRNA target site (red) is shown on top and chromatograms of the same region are shown below. Grey shaded alignments are introns, with allele-discriminating SNPs shown in pink and homologous intron 2 sequences in blue (shifted upstream by 21 bp in *138m1* and *186m2* due to gaps). PAM is underlined and boxed in blue for correspondence with the sequence traces below. Black triangles denote the Cas9 cleavage site and black dashed box corresponds to the 2-bp deletion (−2) detected in KO-5 and KO-69. The two fusion alleles as determined by SNPs are marked below the KO-5 and KO-69 traces (see Supplemental Figure S2 for the full sequence alignment).

### Assessment of off-target activity in mutants

A combination of computational prediction and experimental verification was used to assess off-target effects. Potential off-target sites of the gRNA were predicted by CCTop (Stemmer et al., 2015) using the *P. trichocarpa* v3.1 reference genome as well as the two SNP-substituted Pta717 v2 (*P. alba* and *P. tremula*) subgenomes (Xue et al., 2015). The same four exonic locations were ranked among the top potential off-target sites (excluding intergenic or unassembled scaffold sequences) across the three genomes, each having three mismatches with the gRNA sequence. We designed three sets of primers to examine potential editing at the four off-target sites; OT1 (Potri.004G115600 and Potri.004G118000), OT2 (Potri.004G138000), and OT3 (Potri.014G024400). Amplicon sequencing of 20 trichomeless mutants found no off-target activity across these four sites (Supplemental Dataset S2).

### Absence of triterpenes in trichomeless leaves

Trichomes as epidermal outgrowths are covered with waxy cuticles like other epidermis cells (Hegebarth et al., 2016). The striking glabrous phenotype of the mutants prompted us to compare leaf wax composition between control and trichomeless plants. Total wax load of mature leaves (extractable wax from leaf surface) did not change significantly between genotypes (Figure 5A). Alkanes were the most abundant class of leaf cuticular waxes detected in 717 and differed little between control and trichomeless plants (Figure 5B). In contrast, levels of triterpenes, fatty alcohols and β-sitosterol were significantly reduced in the mutants (Figure 5B-D). Specifically, the wax of mutant leaves was devoid of any triterpenes, including α-amyrin, β-amyrin, β-amyrone and lupenone (Figure 5E). Two primary alcohols, 1-octacosanol (C28) and 1-hexacosanol (C26), were depleted in the mutants by >50% (Figure 5C), and β-sitosterol, by 42% (Figure 5D). To further investigate the absence of triterpenes in the mutant wax, whole leaf tissues were also profiled for compounds that were significantly reduced in cuticular wax. Again, triterpenes were not detected in the leaves of trichomeless mutants (Figure 5E), whereas 1-octacosanol, 1-hexacosanol and β-sitosterol were detected at levels comparable with controls (Figure 5C,D). The data support a previously unsuspected link between triterpene accrual and non-glandular trichomes in poplar.

**Figure 5.**
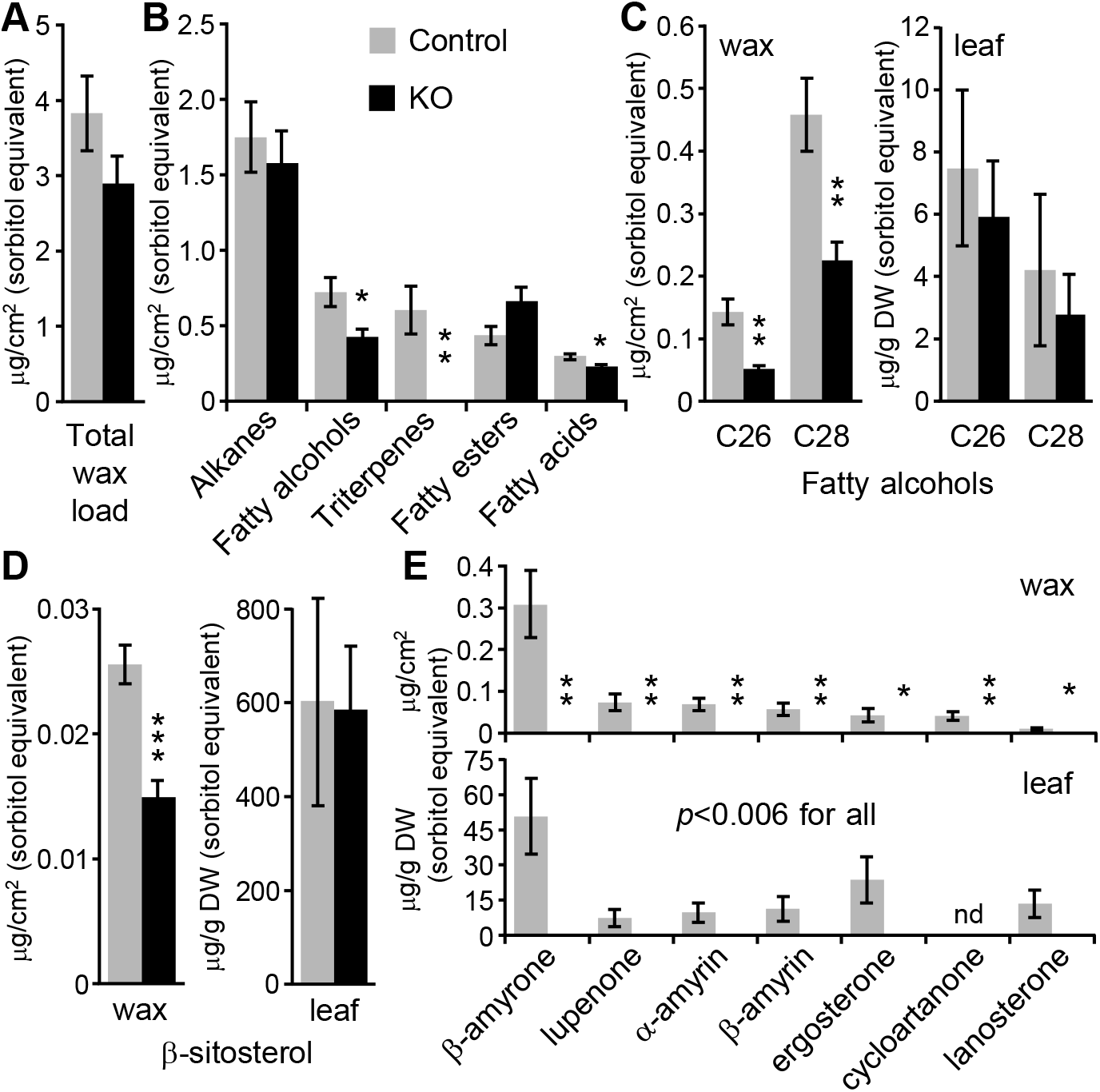
Cuticular wax composition of trichomeless and control leaves. **A**, Total wax load. **B**, Major classes of cuticular wax. **C**, Fatty alcohols (C26, 1-hexacosanol; C28, 1-octacosanol) in wax (left) or whole leaves (right). **D**, β-sitosterol detected in wax (left) or whole leaves (right). **E**, Triterpenes detected in wax (top) or whole leaves (bottom). Ergosterone, 14,24-dimethyl-ergosta-8,25-dien-3-one; cycloartanone, 24-methylene cycloartan-3-one; lanosterone, lanosta-8,24-dien-3-one. Data are mean±SD of n=5. All concentration estimates were based on sorbitol equivalent. Statistical significance was determined by 2-tailed *t*-test (* *P*<0.05, ** *P*<0.01, *** *P*<0.001). nd, not detected.

## DISCUSSION

The present study demonstrates that a single gRNA targeting conserved genomic sites is highly effective for multiplex editing in poplar. The 30 independent KO lines experienced an average of 5.4 CRISPR/Cas9-mediated cleavages per line based on indel alleles, which is likely an underestimate because many NA alleles may also result from CRISPR/Cas9 cleavages as shown for KO-5 and KO-69 (Figures 3 and 4). The unexpected genomic complexity in the hybrid 717 highlights the importance of ensuring SNP-free targets for gRNA design (Zhou et al., 2015), as well as the challenge of decoding multiplexed edits among highly homologous gene duplicates.

The negligible editing by the ΔG construct (Supplementary Dataset S1) provides insight into scaffold structure and stability. The ΔG configuration can lead to two hypothetical outcomes: either the guanine is omitted from the scaffold and the gRNA remains intact and capable of base pairing to the target sites for Cas9 cleavage, or the guanine is sequestered for secondary structure folding of the scaffold, resulting in a 3⍰-truncated gRNA no longer PAM-adjacent at the target sites (Figure 2B). The lack of mutations in ΔG transformants supports the latter scenario and is consistent with transcription and folding of gRNA molecules preceding their base-pairing with genomic targets. Our finding suggests that 3’-truncated gRNA could serve as an alternative approach for generating transgenic controls.

A number of methods are commonly used for decoding CRISPR-mediated mutation patterns, including restriction digestion, endonuclease-based mismatch detection, gene-/allele-specific PCR sometimes in conjunction with cloning and/or Sanger sequencing (Figure 4), or amplicon deep sequencing (Figure 3). The pros and cons of these methods have been discussed elsewhere (e.g., Germini et al., 2018). Analysis of genome editing across multiple target sites poses additional challenges over mono-targeted experiments, especially when highly homologous target and flanking sequences are encountered. These multiplex scenarios generally exceed the resolution of most methods or may require additional assays (*e*.*g*., allele-specific PCR) to determine editing outcomes. In the case exemplified here, deep sequencing of a pooled library of amplicons obtained with consensus primers for all eight target alleles was highly effective for decoding multiplexed edits. The use of consensus primers provides built-in controls for each PCR, allowing for high confidence calling of NA alleles (see Supplemental Dataset S1) which are otherwise difficult to distinguish from failed PCR in individual reactions. As another advantage, the amplicon deep sequencing data can be used for *de novo* assembly which in our case led to the discovery of unexpected copy number variations of MYB186 and MYB138 in the experimental poplar 717 genotype. Although technical limitations remain in short-read mapping to highly homologous sites, inclusion of allele-specific SNPs within the amplicon region and adoption of bioinformatic programs with parameter tuning capabilities (*e*.*g*., AGEseq) are key to multiplexed mutation pattern determination.

The glabrous mutants (Figure 2) provide strong support for an essential role of PtaMYB186/138/38 in the initiation of trichome development in 717. Additionally, the low trichome density of KO-27 suggests that MYB38 plays a redundant but minor role in leaf/stem trichome initiation (Figures 2 and 3). Follow-up research, including allele-specific KOs, is needed to dissect the functional redundancy and allele-dose response of clade 15 MYB members more fully. The unedited (WT) *MYB38* alleles in KO-27 appear stable during vegetative propagation as this event has maintained a low trichome density for over two years in both tissue culture and greenhouse environments. This adds to previously reported stability of CRISPR editing outcomes in clonally propagated poplar (Bewg et al., 2018)

The loss of trichomes did not significantly affect the total epidermal wax load but led to a complete absence of triterpenes both in cuticular wax and whole leaves of the mutants. It is unlikely that MYB186/138/38 have an additional role in triterpene biosynthesis (*i*.*e*., lack of triterpenes as a direct KO effect) because of their recent duplication history (Figure 1) and because a recent report implicated phylogenetically distinct MYBs in triterpene regulation (Falginella et al., 2021). We interpret the absence of triterpenes in trichomeless leaves as suggesting a role for non-glandular trichomes in triterpene accrual in poplar. While glandular trichomes are well known for their roles in biosynthesis and storage of terpenes (Lange and Turner, 2013), the presence of terpenes in non-glandular trichomes has only been reported recently (Santos Tozin et al., 2016; Dmitruk et al., 2019). The genetic evidence presented herein provides strong support for a functional link between triterpenes and non-glandular trichomes that warrants further investigation.

The glabrous phenotype of the null mutants we obtained highlights the potential utility of trichomes as a visual reporter. Assessments of CRISPR/Cas functionality often target the chlorophyll biosynthetic enzyme phytoene desaturase (PDS) (Norris et al., 1995), as mutations result in an albino phenotype (Shan et al., 2013; Ma et al., 2015; Xie et al., 2015). Whilst phenotypically obvious, *PDS* mutations are lethal for the regenerated plant, thus limiting follow-up investigations. Alternatively, the glabrous phenotype achieved by KO of trichome-regulating *MYBs* is non-lethal and no inhibition to plant growth was detected. This allows stacked mutagenesis of these mutants, including reparative transformations to restore trichome initiation. The use of trichomes as a visual reporter for CRISPR/Cas9 mutation or repair of a defective allele has been established in Arabidopsis (Hahn et al., 2017; Hahn et al., 2018) which provides support for further developing this system in poplar.

## MATERIALS AND METHODS

### Generation of KO mutants

The ΔG and KO constructs in p201N-Cas9 (Jacobs et al., 2015) were prepared by Gibson assembly. PCR was used to amplify the p201N-Cas9 binary vector following *Swa*I (New England BioLabs) digestion, and the *Medicago truncatula Mt*U6.6 promoter and scaffold fragments from HindIII and EcoRI (New England BioLabs) digested pUC-gRNA shuttle vector (Jacobs et al., 2015), with Q5 High-Fidelity DNA Polymerase (New England BioLabs) and primers (Sigma) listed in Supplemental Table S1. The p201N-Cas9 (Addgene 59175) and pUC-gRNA (Addgene 47024) plasmids were both gifts from Wayne Parrott. Two pairs of oligos (Sigma) corresponding to the consensus gRNA target site in exon two of *MYB186* (Potri.008G089200), *MYB138* (Potri.008G089700) and *MYB38* (Potri.010G165700) were assembled with p201N-Cas9. The NEBuilder HiFi DNA Assembly Cloning Kit (New England Biolobs) was used to assemble p201N-Cas9, *Mt*U6.6 promoter and scaffold fragments with a pair of oligos containing the gRNA target sequence (Supplemental Table S1). Following transformation into DH5α *E. coli* (Zymo Research Mix & Go! Competent Cells), PCR-positive colonies were used for plasmid purification before Sanger sequencing (Eurofins) confirmation. Plasmids were then heat-shocked into *Agrobacterium tumefaciens* strain C58/GV3101 (pMP90) (Koncz and Schell, 1986) and confirmed by colony PCR.

*Populus tremula* x *alba* (IRNA 717-1B4) transformation and regeneration was performed as outlined in Meilan and Ma (2006), except 0.05 mg/L 6-benzylaminopurine was used in shoot elongation media, and 200 mg/L L-glutamine was added to all media, with 3 g/L gellan gum (PhytoTechnology Lab) as a gelling agent. Following a 2-day agrobacterial cocultivation, leaf discs were washed in sterile water followed by washing in 200 mg/L cefotaxime and 300 mg/L timentin with shaking for 1.5 hr. Transformants were selected on media supplemented with 100 mg/L kanamycin, 200 mg/L cefotaxime and 300 mg/L timentin for callus induction and shoot regeneration and with kanamycin and timentin for shoot elongation and rooting. All cultures were grown and maintained at 22°C under a 16-hr light/8-hr dark photoperiod with Growlite® FPV24 LED (Barron Lighting Group) at ∼150 µmol/m^2^/s.

### RNA-seq analysis

For developmental profiling, LPI-1, LPI-5 and LPI-15 were collected from three greenhouse-grown WT plants (∼5 ft in height) for RNA extraction using Direct-zol RNA MiniPrep kit (Zymo Research) with Plant RNA Purification Reagent (Invitrogen). RNA-seq library preparation and Illumina NextSeq 500 sequencing was performed at the Georgia Genomics and Bioinformatics Core. We obtained 10.8-13.3 PE75 reads per sample. After pre-processing to remove adapter and rRNA sequences, reads were mapped to the 717 SNP-substituted genome sPta717 v2 (Xue et al., 2015) using STAR v2.5.3a (Dobin and Gingeras, 2015). Transcript abundance in FPKM (fragments per kilobase of transcript per million mapped reads) was estimated by featureCounts v1.5.2 (Liao et al., 2014).

### Amplicon sequencing determination of mutation spectrums

Newly emerged leaves were excised from individual events in tissue culture for genomic DNA extraction (Dellaporta et al., 1983). The DNA pellet was resuspended in water with RNase A (10 µg/mL) for amplicon library preparation using GoTaq G2 Green Master Mix (Promega) and primers (Supplemental Table S1) spanning the gRNA target site (between 264 bp to 280 bp). Samples were then barcoded with Illumina amplicon indexing primers and pooled for Illumina MiSeq nano PE150 sequencing performed at the University of Georgia’s Georgia Genomics and Bioinformatics Core. Demultiplexed sequence reads were analyzed by the AGEseq (Analysis of Genome Editing by Sequencing) program (Xue and Tsai, 2015), with mismatch allowance set at 1%, followed by manual curation.

Because initial amplicon data analysis revealed lower editing efficiencies (<90%) than we typically observed in 717 (Zhou et al., 2015; Bewg et al., 2018) at several target sites, we performed *de novo* assembly of WT amplicon reads using Geneious, and recovered seven distinct alleles. We then searched the JGI draft 717 genome assembly v1.0 with the *P. trichocarpa* Nisqually-1 v3.1 (Phytozome v12) *MYB186, MYB138* and *MYB38* gene models and extracted the surrounding 50-150 Kb regions from Chr8 and Chr10 for manual annotation against the *P. trichocarpa* Nisqually-1 reference (Figure 3A). The matching *MYB* gene sequences were extracted for error correction using 717 resequencing data (Xue et al., 2015). Curated sequences were used for new (amplicon and allele-/gene-specific) primer design and as references in amplicon data analysis. In the case of WT and transgenic controls with no editing, erroneous read assignments—and hence indel calls—still remained because the amplicon region between some alleles differs only in the number of intronic dinucleotide (GT) repeats (Supplemental Dataset S1). Misassigned reads led to erroneous indel calls of -2, +2 or their multiples outside of the gRNA target site. For this reason, WT and control samples were processed by ustacks from Stacks 2.3 (Catchen et al., 2011). Parameters were adjusted to avoid collapsing reads with SNPs and/or Indels from paralogous alleles into the same tag group and gapped alignments were disabled. Tags from the output were then used for allele assignment.

### Determination of leaf and cuticle wax compositions

Leaf punches (25 mm diameter) were taken from mature leaves of similar size (between LPI-10 and LPI-15) of soil-grown plants in a growth chamber and washed in 4 mL of methylene chloride for 30 sec. The washes were dried under a continuous N_2_ stream before resuspension in 400 µL chloroform. A 200 µL aliquot was subsequently dried under vacuum and the residues shipped to the Oak Ridge National Laboratory for analysis. Sorbitol (1 mg/mL) was added to the residues as an internal standard and re-dried under N_2_. For whole leaf analysis, liquid nitrogen-ground and freeze-dried powders from LPI-5 (25 mg) of control and KO plants were extracted by 80% ethanol to which sorbitol (1 mg/mL) was added and dried under N_2_. The samples were derivatized prior to analysis on an Agilent Technologies 7890A GC coupled to a 5975C inert XL MS fitted with an Rtx-5MS capillary column with a 5m Integra-Guard column (Restek) as described in Holwerda et al. (2014). Compound identification was based on mass spectral fragmentation patterns against the NIST08 database and an in-house library built with authentic standards.

### Tissue Imaging and SEM analysis

Images of poplar were taken with either a Google Pixel 3a running Android v11, or a Leica M165 FC dissection microscope attached to a Leica DFC500 camera running Leica Application Suite software v3.8.0. Scanning electron microscopic (SEM) observations were obtained using Hitachi 3400 NII (Hitachi High Technologies America) microscope following optimized protocols at the Center for Ultrastructural Research at the Fort Valley State University. LPI-1 from growth chamber plants or young leaves of tissue culture plants were processed for primary fixation at 25°C in 2 % glutaraldehyde (Electron Microscopy Sciences, EMS) prepared with Sorensen’s Phosphate buffer, pH 7.2 (EMS) for one hour and then washed three times for 15 min each with the same buffer before secondary fixation in 1% osmium tetroxide (EMS) prepared in Sorensen’s Phosphate buffer, pH 7.2 for 1 hour at 25°C. After three washes with dH_2_O for 15 min each, fixed tissues were dehydrated with ethanol series passing through 25%, 50%, 75%, and 95% for 15 min each, followed by three changes of 100% ethanol for 15 min each. Critical point drying of fixed samples was conducted using a critical point dryer (Leica) and then samples were placed on Hitachi M4 aluminum specimen mounts (Ted Pella) using double sided carbon adhesive tabs (EMS) for coating. Gold coating of 50 Å thickness was done for 60 sec using sputter coater (Denton Desk V) under a vacuum pressure of 0.05 torr. Image acquisition in various magnification was done at accelerating voltage of 5 KV.

## ACCESSION NUMBERS

The RNA-seq data has been deposited to the National Center for Biotechnology Information’s Sequence Read Archive under accession No. PRJNA753499.

## ACKNOWLEDGEMENTS

The authors thank Gilles Pilate of the Institut National de la Recherche Agronomique, France for providing poplar clone INRA 717-1B4, Hongduyen Pham and Margot Chen for tissue culture assistance, Yingying Zhu for RNA from developmentally staged leaves and Liang-Jiao Xue for guidance on RNA-seq data processing. We additionally thank the Department of Energy Joint Genome Institute and collaborators for prepublication access to the *Populus tremula* × *P. alba* (IRNA 717-1B4) genome sequence and annotation.

## SUPPORTING INFORMATION

**Table S1**. Primers used in this study.

**Figure S1**. PCR confirmation of NA alleles using allele-specific primers.

**Figure S2**. Sequence alignment of wild type and fusion *MYB* alleles from KO-5 and KO-69.

**Dataset S1**. CRISPR/Cas9 mutation patterns of the eight target *MYB* alleles in ΔG and KO lines. **Dataset S2**. Assessment of off-target activity in trichomeless mutants.

